# A low-cost *rpoB*-based multiplex MAMA PCR for differentiation of the *Klebsiella pneumoniae* species complex

**DOI:** 10.64898/2026.04.14.718422

**Authors:** Mehbooba Sharmin, Al Amin, Hafizur Rahman, Nicol Janecko, Samir K Saha, Yogesh Hooda, Arif Mohammad Tanmoy, Senjuti Saha

**Author notes:** Correspondence to Senjuti Saha, PhD, MPH, Deputy Executive Director, Child Health Research Foundation, 23/2 Khilji Road, Shyamoli, Dhaka 1206, Bangladesh.

## Abstract

The *Klebsiella pneumoniae* species complex (KpSC) is a clinically important group of closely related pathogens associated with invasive infections. The complex comprises seven closely related members, which are often reported as *K. pneumoniae*, particularly in resource-limited settings. Accurate differentiation of KpSC members remains challenging because routine laboratory methods lack sufficient resolution, and approaches like mass spectrometry and whole genome sequencing (WGS) are not widely available. Consequently, the epidemiology and clinical significance of non-*K. pneumoniae* members of the KpSC remain underrecognized. We developed a conventional multiplex mismatch amplification mutation assay (MAMA) PCR targeting species- and subspecies-specific single-nucleotide polymorphisms in the housekeeping gene *rpoB*, with six primer sets for differentiation of common KpSC members. The assay was validated against 49 genomically characterized clinical isolates, after which 179 wastewater-derived isolates provisionally identified as *Klebsiella* spp. by standard microbiological methods were tested. Of these, 174 were assigned to specific KpSC members by the assay, while 5 produced inconclusive amplification patterns. A subset of 16 environmental isolates was selected for WGS, including four of the five inconclusive isolates. All environmental isolates with interpretable MAMA PCR patterns were concordant with WGS. The four inconclusive environmental isolates were identified as *Enterobacter* spp. Overall, comparison of MAMA PCR with WGS showed 100% sensitivity and 100% specificity for all tested targets, and the total cost was approximately US$1. This *rpoB*-based multiplex MAMA PCR provides a simple, accurate, and low-cost approach for differentiation of KpSC members in routine laboratories and may support improved identification and surveillance in resource-limited settings.

**Importance:** The *Klebsiella pneumoniae* species complex (KpSC) has seven members but is often reported as a single organism in routine laboratories, masking clinically and epidemiologically important diversity. As a result, the contribution of non-*K. pneumoniae* KpSC members to human and environmental microbiology remains poorly defined, especially in low-resource settings. We developed a conventional multiplex mismatch amplification mutation assay (MAMA) PCR based on discriminatory *rpoB* single nucleotide polymorphisms for differentiation of common KpSC members using standard PCR and agarose gel electrophoresis. The assay demonstrated 100% sensitivity and 100% specificity against whole-genome sequencing and excluded non-*Klebsiella* environmental isolates initially identified as *Klebsiella pneumoniae* using standard microbiological procedures. With an estimated per-test cost of about US$1, this method offers an affordable and scalable option for laboratories seeking more accurate KpSC identification and improved surveillance.

## Introduction

*The Klebsiella pneumoniae* species complex (KpSC) is a clinically important group of closely related Gram-negative bacteria associated with bloodstream, respiratory, and urinary tract infections and other invasive infections. One member of KpSC, *Klebsiella pneumoniae*, is recognized as a World Health Organization priority pathogen because of increasing carbapenem resistance (1, 2). Global burden analyses indicate that South Asia bears a high burden of *K. pneumoniae* infection, characterized by elevated mortality, particularly among neonates and children under five in lower sociodemographic settings (3). In Bangladesh, it contributes substantially to neonatal sepsis and antimicrobial resistance associated disease burden, highlighting the need for accurate identification and surveillance (4).

However, genomic studies have shown that many isolates identified as *K. pneumoniae* by conventional methods actually belong to KpSC, a group of closely related species and subspecies with 96% to 97% average nucleotide identity that differ in epidemiology, antimicrobial resistance carriage, virulence factor carriage, and clinical outcomes (2, 5–8). KpSC includes *Klebsiella pneumoniae* (Kp1), *Klebsiella quasipneumoniae* subsp. *quasipneumoniae* (Kp2), *Klebsiella variicola* subsp. *variicola* (Kp3), *Klebsiella quasipneumoniae* subsp. *similipneumoniae* (Kp4), *Klebsiella variicola* subsp. *tropica* (Kp5), *Klebsiella quasivariicola* (Kp6), and *Klebsiella africana* (Kp7) (2, 6). Beyond *K. pneumoniae*, species such as *K. variicola* and *K. quasipneumoniae* are increasingly recognized as clinically relevant pathogens with distinct host associations and disease patterns (5, 8). This is particularly important in high burden settings such as Bangladesh, where non *K. pneumoniae* KpSC members can make up to 15% of the isolates recovered from neonates, but are neglected because of limited tools to identify and document their epidemiology (4).

Accurate identification of KpSC members remains challenging because currently available diagnostic approaches do not adequately balance resolution, accessibility, and long-term reliability (9). Conventional phenotypic methods and 16S rRNA sequencing lack sufficient resolution to distinguish closely related members of the complex, whereas higher resolution approaches such as MALDI TOF mass spectrometry and whole genome sequencing (WGS) require specialized infrastructure that are not routinely accessible in many low- and middle-income settings (10–12). PCR-based methods offer a more accessible alternative and have improved detection of major species. However, existing assays do not cover all clinically relevant or common members of the complex, do not achieve subspecies level resolution, or rely on nonconserved or accessory targets that may limit long term robustness (13–17). This gap is especially consequential in high-burden settings, where misidentification obscures surveillance, underestimates the contribution of non-K. *pneumoniae* KpSC members, and limits accurate tracking of antimicrobial resistance and disease burden. There is therefore a clear need for a simple, low-cost, and robust method that can differentiate KpSC members in routine laboratory settings.

Housekeeping genes of KpSC provide an optimal target because they are essential, conserved under stabilizing selection, and still contain sufficient sequence variation to discriminate closely related taxa (18). Among these, the RNA polymerase β subunit gene, *rpoB*, has been widely recognized as a robust marker for bacterial identification, offering adequate resolution within the KpSC (7, 19). Discriminatory single-nucleotide polymorphisms (SNPs) within conserved regions can therefore be leveraged for simple molecular identification without reliance on sequencing-based methods. The mismatch amplification mutation assay (MAMA) exploits such SNPs through allele specific primer design and has previously been used for rapid detection of clinically relevant closely-related pathogens, including azithromycin resistance associated mutations in *Salmonella* (20).

In this study, we aimed to develop a low-cost conventional multiplex MAMA PCR for species and subspecies level identification within the KpSC. Specifically, we identified discriminatory SNPs within *rpoB*, designed a multiplex MAMA PCR targeting these SNPs, validated the assay using clinical isolates representing major circulating sequence types in Bangladesh, and evaluated its applicability using environmental isolates.

## Methods

### Identification of discriminatory *rpoB* SNPs within the KpSC

To evaluate the suitability of *rpoB* as a discriminatory target within the KpSC, a total of 1,082 genomes representing six phylogroups of the KpSC were retrieved from NCBI GenBank (accessed 23 August 2025); Kp5 was not included because of unavailability of sequenced genomes from clinical sources. Of these, 982 genomes were selected by restricting the dataset to RefSeq annotated genomes (GCF accessions) and excluding contig level assemblies to ensure consistent annotation quality. Only scaffold level or higher quality assemblies were retained for analysis. In addition, 100 genomes from a previously described *K. pneumoniae* diversity panel were included to capture a broad range of genetic diversity (21).

*rpoB* sequences were extracted from each genome using a custom BLAST based pipeline implemented in bash. Briefly, each genome assembly was formatted as a nucleotide database, and the 4,029 bp *rpoB* sequence from *K. pneumoniae* reference strain HS11286 (GenBank accession GCF_000240185.1) was used as the query for local BLASTN searches with outfmt 6 output. Matching regions were extracted on the basis of alignment coordinates and manually inspected to confirm successful retrieval of the *rpoB* locus. The full extraction script is provided in the Supplementary Materials (Data S1).

Extracted *rpoB* sequences were aligned using MAFFT v7.526 (22). A maximum likelihood phylogeny was then inferred using IQ TREE 2 v3.0.1 (23), and the resulting tree was visualized and annotated in iTOL v7 (24). Details of all 1,082 genomes, including accession numbers and species assignments, are provided in Table S1.

Discriminatory single-nucleotide polymorphisms (SNPs) within *rpoB* were identified by manual inspection of the multiple sequence alignment in MEGA v12.1 (25). SNPs were considered species discriminatory if they were conserved in all genomes of a given species and absent from all other KpSC members. SNP positions were based on reference genomes, except for *K. quasipneumoniae* subsp. *quasi*pneumoniae and subsp. *similipneumoniae*, for which representative RefSeq-annotated and ATCC strains were used respectively. For subspecies level differentiation within *K. quasipneumoniae*, SNPs were selected if they were conserved in all genomes of subsp. *quasipneumoniae* and absent from all genomes of subsp. *similipneumoniae*. These discriminatory SNPs were subsequently used for primer design.

### Primer design

Regions within the *rpoB* gene were selected based on the presence of discriminatory SNPs identified from sequence alignment and conserved flanking sequences suitable for primer design. The selected loci were first assessed for specificity using online BLASTN (https://blast.ncbi.nlm.nih.gov/Blast.cgi) against the NCBI nucleotide database (nt). Regions showing high specificity with no significant off-target matches were selected. Primer properties, including length, melting temperature (Tm), GC content, secondary structures, primer-dimer formation were evaluated using the OligoAnalyzer web tool (https://www.idtdna.com/pages/tools/oligoanalyzer) (Integrated DNA Technologies, USA).

A common forward primer was designed from a conserved region across all members of the KpSC, enabling binding to all members of the complex. The forward primer position was selected to generate distinct amplicon sizes when paired with different reverse primers, enabling clear target differentiation by agarose gel electrophoresis. Species and subspecies-specific reverse primers were designed based on discriminatory SNPs to develop MAMA PCR. For each target, the SNP was positioned near the 3′ end of the reverse primer (within the last three nucleotides) to ensure specificity. An additional mismatch was introduced either at the penultimate position or at the third nucleotide from the 3’ end to further enhance discrimination, following established MAMA PCR design principles (26, 27). An internal PCR control was designed using the common forward primer and an additional reverse primer targeting a conserved region of the *rpoB* gene shared across all KpSC members. This control primer was not SNP-specific and was included to confirm successful amplification across all members of the complex.

Apart from the control reverse primer (KpSc_rpoB_1036_Rev), the assay consisted of one common forward primer and six target-specific reverse primers, each set theoretically generating distinct amplicon sizes. A two reaction PCR system was implemented to accommodate primer compatibility and amplicon separation: PCR-1 targeted *K. pneumoniae, K. quasipneumoniae* subsp. *quasipneumoniae, K. quasipneumoniae* subsp. *similipneumoniae* and *K. variicola*, while PCR-2 targeted *K. africana* and *K. quasivariicola*. All primers were synthesized commercially by Macrogen (Seoul, South Korea) and Integrated DNA Technologies (IDT, Coralville, IA, USA).

### PCR optimization

For initial optimization of the multiplex MAMA PCR, 8 KpSC isolates were selected from a prior study based on their species and subspecies identity to ensure balanced representation, including *K. pneumoniae* (n = 2), *K. quasipneumoniae* subsp. *quasipneumoniae* (n = 2), *K. quasipneumoniae* subsp. *similipneumoniae* (n = 2) and *K. variicola* (n = 2) (4), Table S2. Briefly, isolates were originally cultured from clinical samples collected from pediatric patients in Bangladesh with invasive infections, including blood and cerebrospinal fluid. Standard culture-based methods were used, and isolates were characterized using biochemical tests: Triple Sugar Iron (TSI, Oxoid, Basingstoke, UK), Motility Indole Urease (MIU, HIMEDIA, Mumbai, India), and Simmon’s Citrate Agar (SCA, Oxoid, Basingstoke, UK) following standard microbiological procedures. Isolates showing an acidic slant and butt in TSI (A/A), non-motility in MIU, indole-negative, urease activity (variable) and citrate-positive profiles were identified as *Klebsiella pneumoniae*. Isolates were preserved in aliquots of skim milk, tryptone, glucose, glycerol (STGG) medium and stored at −80°C. To confirm their identity at the species and subspecies levels, WGS was performed on an in-house Illumina NextSeq 2000 platform (Illumina, San Diego, CA, USA) with 150 bp paired end reads.

Template DNA was prepared using the boiling lysis method. A single colony from an overnight culture of the isolate on MacConkey agar (Oxoid, UK) was suspended in 60 µL of TE buffer. The suspension was incubated at 95 °C for 15 min, then centrifuged at 7000 rpm for 5 min. The resulting supernatant was used as the DNA template for downstream PCR assays.

Conventional PCR reactions were performed using a thermal cycler (ProFlex PCR Systems; Thermo Fisher Scientific, Waltham, MA, USA) and HOT FIREPol mastermix (Cat. No: 04-27-00115; Solis BioDyne, Tartu, Estonia) according to the manufacturers’ protocols. The performance of the assay was initially evaluated using singleplex PCR reactions, followed by duplex and triplex configurations, and finally optimized as a multiplex PCR assay. The reaction volume was set to 25 µL, and 4% DMSO (Dimethyl sulfoxide) was added to PCR-1 reactions to reduce secondary structures and improve multiplex amplification of GC-rich regions.

The PCR cycling conditions were as follows: initial denaturation at 95°C for 15 minutes (1 cycle), followed by 34 cycles of denaturation at 95°C for 30 seconds, annealing at 65°C for 30 seconds, and extension at 72°C for 40 seconds, with a final extension at 72°C for 3 minutes (1 cycle) and a hold at 4°C. Details of the PCR assay, including primer sequences, concentration and target species are summarized in Table 1. Following PCR, amplified products were run on 1.2% agarose gels at 110 V for 75 min and visualized using a Gel Doc XR+ system (Bio-Rad, Richmond, CA, USA).

**Table 1.**
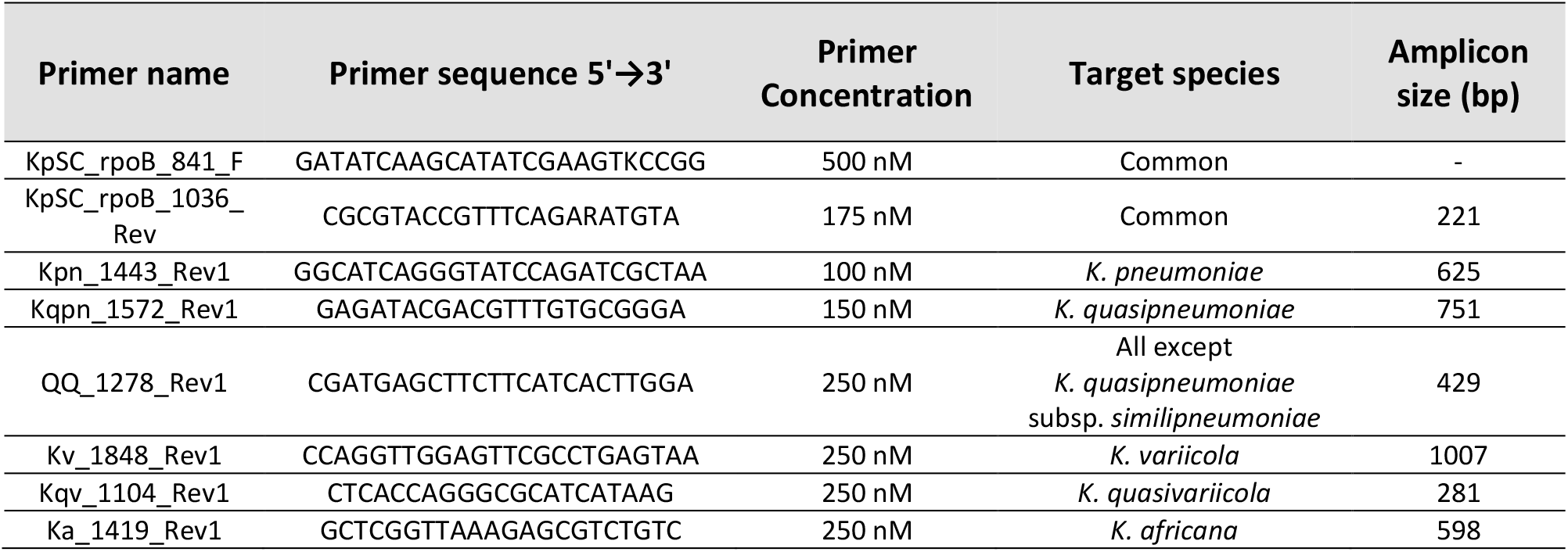
Primer details, including sequences, concentrations, target species and amplicon sizes for multiplex MAMA PCR targeting the *Klebsiella pneumoniae* species complex (KpSC)

### Validation of the MAMA PCR

Further validation was performed using a set of 49 invasive clinical KpSC isolates that were also sequenced in the prior study described above (4). This set included isolates representing the 15 most frequently found sequence types of *K. pneumoniae* in Bangladesh (4). In addition, isolates of *K. quasipneumoniae* subsp. *quasipneumoniae* (n=15) and subsp. *similipneumoniae* (n=15) and *K. variicola* strains (n=4) were included, reflecting their availability in the dataset. Each isolate represented a unique sequence type (ST). A complete list of the strains and their corresponding STs is provided in Table S2.

### Isolation and identification of environmental *Klebsiella* strains

A total of 140 wastewater samples were collected in August 2023 from seven administrative areas of Dhaka, which include Adabor, Kafrul, Pallabi, Mirpur, Mohammadpur, Sher-e-Bangla Nagar, and Shah Ali (Fig. S1). Sampling sites were selected to minimize cross-contamination and repeated collection, maintaining a median distance of 0.44 km between locations. Approximately 10 mL of wastewater was collected from each site using a sterile 15 mL conical tube attached to a rope. The samples were then transferred into new sterile tubes under aseptic conditions and transported to the laboratory.

Following collection, samples were centrifuged at low speed (1000 rpm for 1 minute) to remove large debris, and the supernatant was transferred to a fresh tube. Serial dilutions (10^−1^ to 10^−3^) were prepared using 0.9% sterile saline. A volume of 100 µL from the 10^−2^ and 10^−3^ dilutions were spread plated onto MacConkey agar plates and incubated overnight at 37 °C. After incubation, colonies displaying *Klebsiella*-like morphology (large, mucoid, lactose-fermenting, pink-colored colonies) were selected, and three colonies per sample were sub-cultured onto fresh MacConkey agar plates to obtain pure isolates.

All selected isolates were subjected to biochemical characterization as described above using TSI, MIU and SCA tests. After biochemical identification as *Klebsiella pneumoniae*, all environmental isolates were tested using the MAMA PCR-1. Based on the amplification patterns, isolates from each observed phylogroup were selected for WGS, with three isolates randomly chosen per phylogroup. In addition, four out of five isolates showing unusual or unclear amplification patterns were selected based on their distinct banding pattern and were also included to ensure a broader assessment of the assay.

### Whole genome sequencing and bioinformatic analysis

All selected strains (n=16) for WGS were freshly cultured on MacConkey agar plates from STGG stock and incubated overnight (∼16–18 h) at 37 °C. A single colony was picked and subcultured on MacConkey agar to obtain a pure culture. A suspension was then prepared from the overnight culture in 800 µL of 1× PBS, and genomic DNA was extracted using the TIANamp Bacteria DNA Kit (Cat. no. GDP302-02, TIANGEN Biotech, China). Sequencing libraries were prepared using NEBNext Ultra II FS DNA Library kit (Cat: E7805, New England 470 Biolabs, Ipswich, MA, USA) and the Twist Library Preparation EF Kit 2.0 (Cat. No. 104207, Twist Bioscience, South San Francisco, CA, USA) according to the manufacturer’s instructions. Libraries were sequenced on an Illumina NextSeq 2000 platform (Illumina, San Diego, CA, USA) in Dhaka, Bangladesh using 150 bp paired end reads.

Raw sequence data quality was assessed using FastQC v0.11.9. (https://www.bioinformatics.babraham.ac.uk/projects/fastqc/) Adapter trimming and quality filtering (phred score= 20) were performed using fastp v0.23.4 (28). *De novo* genome assembly was conducted using Unicycler v0.5.1 (default parameters) (29). Species identification was performed using the Pathogenwatch web platform (https://pathogen.watch/) (30). All fastq files of all isolates sequenced for this study have been submitted to ENA under project accession ERP173558 (PRJEB90555).

### Statistical Analysis and Cost Calculation

The sensitivity and specificity testing was conducted using R version 4.2.2 and the *epiR* package. The cost of the multiplex MAMA-PCR was estimated based on the local prices of consumables used per reaction, including PCR master mix, nuclease-free water and primers. In addition, local costs of reagents used for gel electrophoresis, including agarose powder, SYBR Safe stain, and DNA ladder were also considered. As the assay requires two PCR reactions, the total cost per sample reflects the combined cost of both reactions.

## Results

### Identification of discriminatory *rpoB* SNPs and multiplex assay design

Analysis of the *rpoB* gene across the selected 1082 KpSC genomes showed distinct clustering at both the species and subspecies levels. The maximum likelihood phylogeny resolved the complex into distinct clades corresponding to *K. pneumoniae, K. quasipneumoniae, K. variicola, K. quasivariicola*, and *K. africana*. Within *K. quasipneumoniae*, the two subspecies, subsp. *quasipneumoniae* and subsp. *similipneumoniae*, formed well defined subclades (Fig. 1). Overall, genomes belonging to the same species clustered together with clear separation from other members of the complex, supporting *rpoB* as a discriminatory target for assay development.

**Figure 1.**
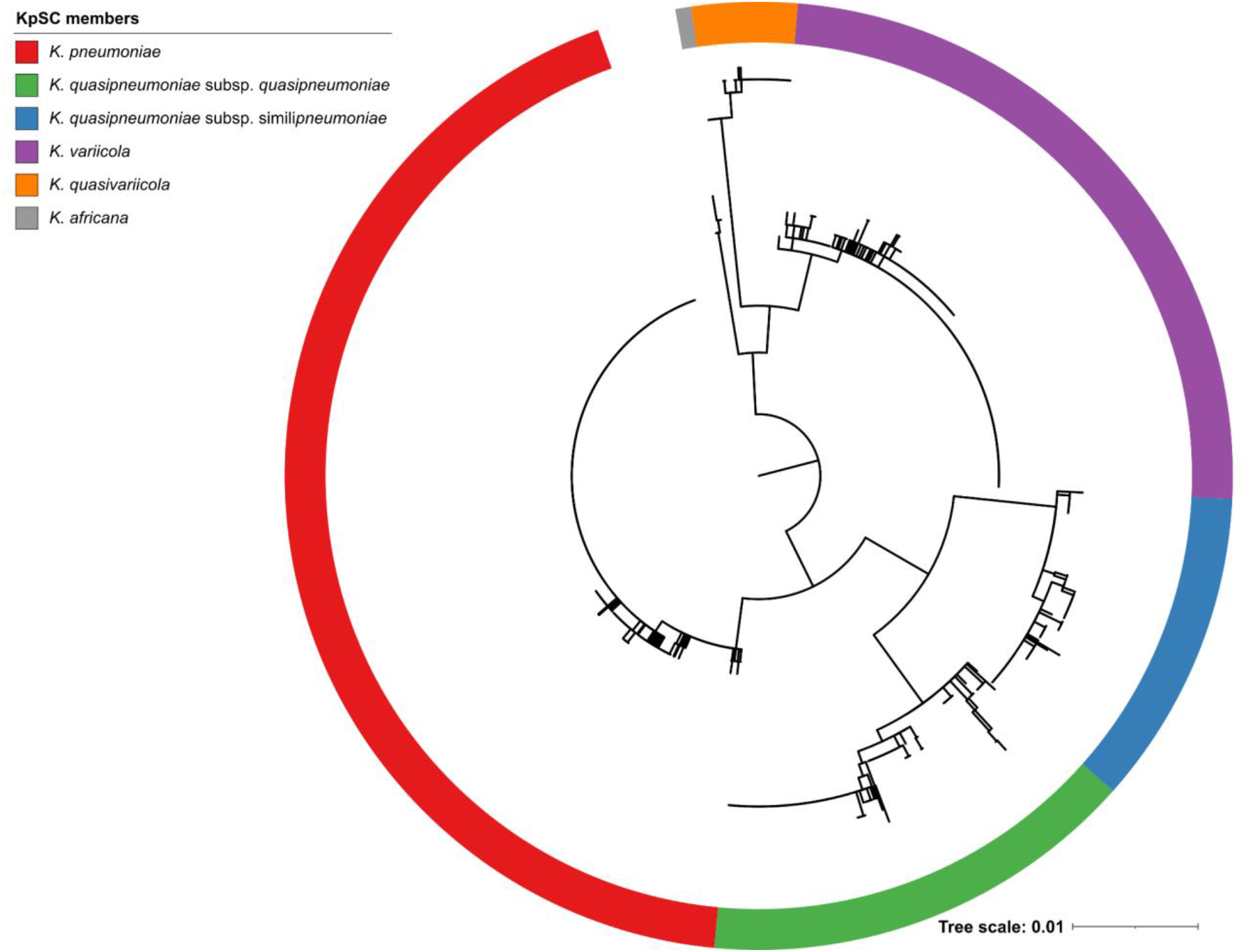
Maximum likelihood tree demonstrating the phylogenetic relationship among members of the *Klebsiella pneumoniae* species complex (KpSC) based on *rpoB* gene sequences. A midpoint rooted phylogenetic tree was constructed using *rpoB* sequences from 1082 genomes described in the legend, with the species indicated by colour in the outer ring.

Comparative analysis of the *rpoB* alignment identified multiple candidate SNPs that were conserved within individual KpSC members and distinct from the remaining members of the complex. From these, discriminatory SNPs were selected for primer design at nucleotide positions 1443 for *K. pneumoniae*, 1572 for *K. quasipneumoniae*, 1848 for *K. variicola*, 1104 for *K. quasivariicola*, and 1419 for *K. africana* (Fig. 2). For *K. quasipneumoniae*, subspecies differentiation required a two-step logic: a species-specific SNP at position 1572 to identify *K. quasipneumoniae*, together with an additional SNP at position 1278 to distinguish subsp. *quasipneumoniae* from subsp. *similipneumoniae*. These SNPs were incorporated into a multiplex MAMA PCR using one common forward primer, six target specific reverse primers, and an internal control primer pair designed to amplify all KpSC members (Fig. 3A). An additional mismatch was introduced within the 3′ terminal region (within the last three bases), positioned either upstream or downstream of the SNP depending on the target.

**Figure 2.**
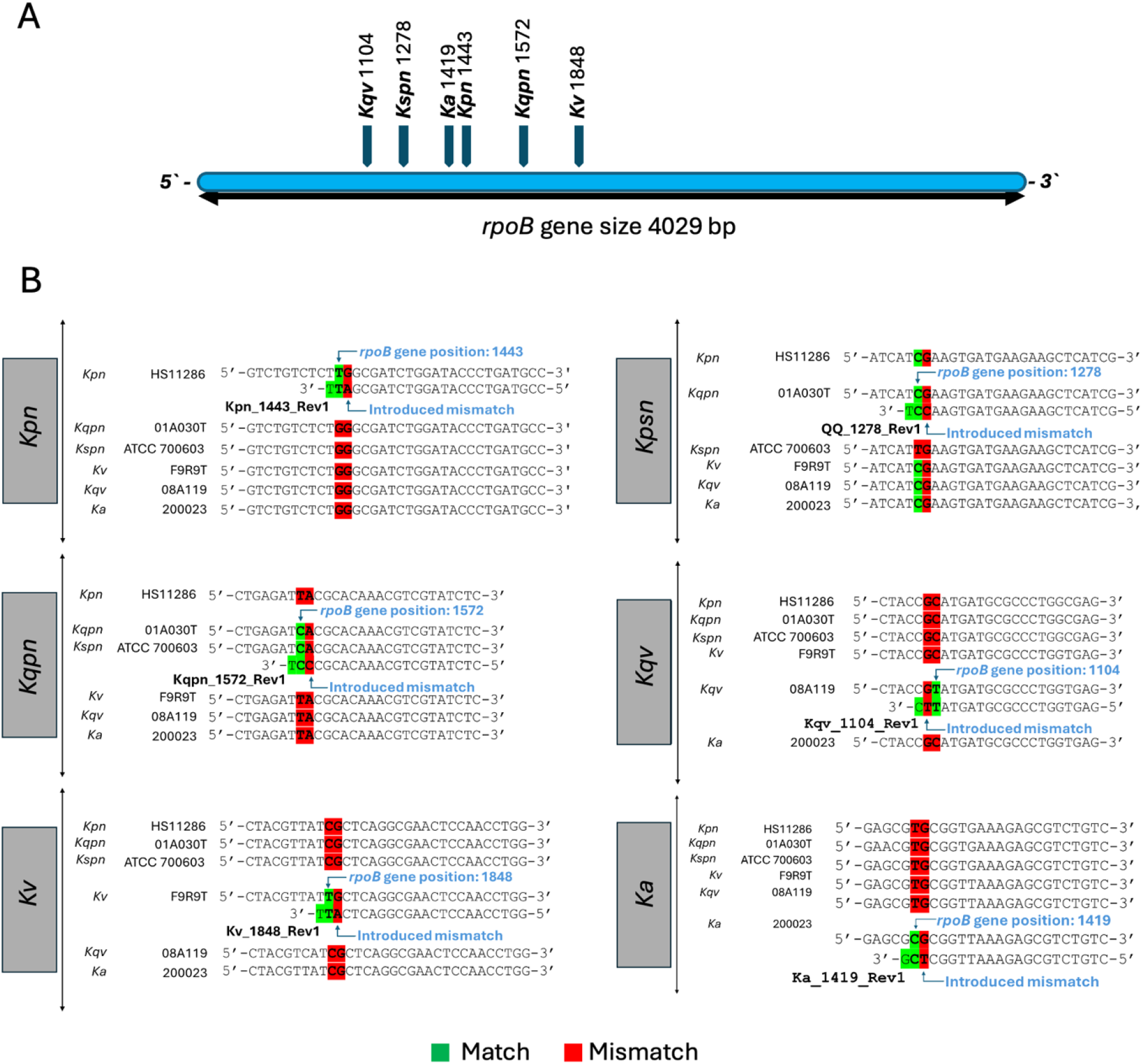
Identification of discriminatory SNPs in the *rpoB* gene and MAMA PCR primer design strategy for differentiation of the *Klebsiella pneumoniae* species complex (KpSC). (A) Schematic representation of the *rpoB* gene (4,029 bp) showing the positions of species and subspecies-specific SNPs used for differentiation of KpSC members. Arrows indicate nucleotide positions of the discriminatory SNPs. (B) Alignment-based representation of reverse MAMA primer binding regions for each target species and subspecies. Each panel shows target and non-target species and subspecies grouped to illustrate primer design strategy. Strain names are indicated on the left. Reverse MAMA primers are shown aligned to their target regions. The discriminatory SNP is positioned within the last three nucleotides at the 3′ end of the primer. The green highlights indicate nucleotide matches between primer and template and the red highlights indicate mismatches, including both the discriminatory SNP and the introduced mismatch. [Kpn = *Klebsiella pneumoniae*; Kqpn = *Klebsiella quasipneumoniae* subsp. *quasipneumoniae*; Kspn = *Klebsiella quasipneumoniae* subsp. *similipneumoniae*; Kv = *Klebsiella variicola*; Kqv = *Klebsiella quasivariicola*; Ka = *Klebsiella africana*.]

**Figure 3.**
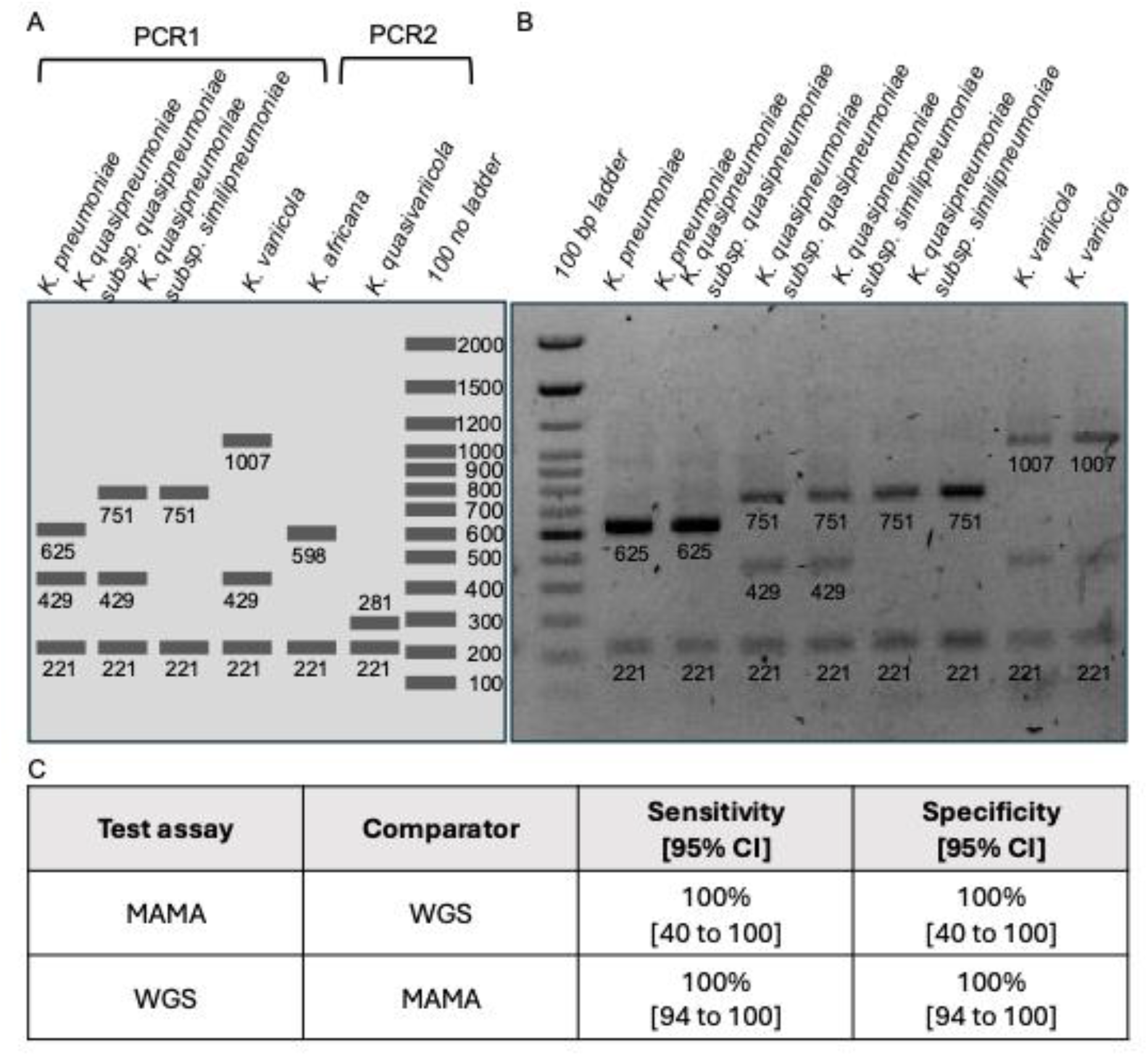
Design and validation of the two-reaction multiplex MAMA PCR assay for differentiation of members of the *Klebsiella pneumoniae* species complex (KpSC). (A) Schematic representation of expected amplification patterns for each target species and subspecies, showing species-specific amplicon sizes alongside a common internal control band. (B) Agarose gel electrophoresis showing amplification patterns obtained using the multiplex MAMA PCR assay (PCR-1) with one common forward primer and five reverse primers. The accompanying table (on the right) summarizes primer names, target specificity and expected amplicon sizes. Species-specific bands were observed at the expected sizes, with each target generating a distinct banding pattern. A common internal control band (221 bp) was present across all lanes. A 100 bp DNA ladder was loaded on both sides of the gel as a molecular size marker. (C) Performance of the assay compared with whole genome sequencing (WGS), showing 100% sensitivity and specificity with corresponding 95% confidence intervals.

### Optimization and validation of the multiplex MAMA PCR

The final assay was optimized as a two-reaction system, designated PCR 1 and PCR 2, with a maximum of three target specific bands detectable in a single reaction. Under the optimized conditions, PCR 1 produced clear and reproducible banding patterns for *K. pneumoniae, K. quasipneumoniae* subsp. *quasipneumoniae, K. quasipneumoniae* subsp. *similipneumoniae*, and *K. variicola* when resolved by agarose gel electrophoresis (Fig. 3B, Fig. S2). The internal control band at 221 bp was consistently detected across KpSC isolates, confirming amplification of members of the complex under the optimized assay conditions. PCR 2 for *K*.*quasivariicola* and *K*.*africana* was not conducted experimentally as confirmed isolates of these species are rare and not available in Bangladesh.

### Validation of the assay using clinical isolates

Following initial optimization, the final multiplex MAMA PCR assay was validated using 49 clinical isolates representing members of the KpSC, all with prior WGS-based species assignments (4) (Table S2). The assay correctly identified *K. pneumoniae, K. quasipneumoniae*, and *K. variicola* at the species level, and further resolved *K. quasipneumoniae* into subsp. *quasipneumoniae* and subsp. *similipneumoniae*. All results were fully concordant with WGS-based classification, corresponding to 100% sensitivity across the 49 clinical isolates tested (Fig. S3).

### Evaluation of the assay using environmental isolates

A total of 179 isolates were recovered from 140 wastewater samples (Fig. S1) and were provisionally identified as *Klebsiella* species by standard microbiological and biochemical methods. All 179 isolates were then screened using the multiplex MAMA PCR for identification within the KpSC (Fig. S4A).

Of these, 135 (75%) were identified as *K. pneumoniae*, 22 (12%) as *K. quasipneumoniae* subsp. *similipneumoniae*, 5 (3%) as *K. quasipneumoniae* subsp. *quasipneumoniae* and 12 as (7%) *K. variicola*. The remaining five (3%) isolates produced inconclusive amplification patterns (Fig. S4B). Although additional bands were observed in two recurring patterns, these results could not be interpreted reliably in the absence of the control band and were therefore considered non-interpretable and not a member of KpSC.

To further assess the environmental isolate classifications, a subset of 16 isolates was selected for WGS, including 12 isolates with interpretable MAMA PCR patterns and 4 of the 5 isolates with inconclusive amplification patterns (Table 2). Selected isolates were sequenced and species assignments were determined from assembled genomes using Pathogenwatch (30). All 12 (12/12, 100%) environmental isolates with interpretable MAMA PCR patterns were fully concordant with WGS, confirming species and subspecies assignments for *K. pneumoniae, K. quasipneumoniae* subsp. *quasipneumoniae* and subsp. *similipneumoniae*, and *K. variicola*. The 4 sequenced isolates with inconclusive MAMA PCR patterns were all identified as members of the genus *Enterobacter*, including 2 *E. cloacae*, 1 *E. roggenkampii*, and 1 *E. hormaechei*.

**Table 2.** Comparison of biochemical identification, MAMA PCR results and WGS-based classification of selected isolates.

| Sequence ID | Biochemical result | MAMA - PCR result |  | WGS result |  |
| --- | --- | --- | --- | --- | --- |
|  |  | Species | Subspecies | Species | Subspecies |
| CHRF_ENV_04 | *Klebsiella* | unidentified |  | *Enterobacter roggenkampii* |  |
| CHRF_ENV_05 | *Klebsiella* | unidentified |  | *Enterobacter hormaechei* |  |
| CHRF_ENV_06 | *Klebsiella* | unidentified |  | *Enterobacter cloacae* |  |
| CHRF_ENV_07 | *Klebsiella* | unidentified |  | *Enterobacter cloacae* |  |
| CHRF_ENV_01 | *Klebsiella* | *K. pneumoniae* |  | *K. pneumoniae* |  |
| CHRF_ENV_02 | *Klebsiella* | *K. pneumoniae* |  | *K. pneumoniae* |  |
| CHRF_ENV_03 | *Klebsiella* | *K. pneumoniae* |  | *K. pneumoniae* |  |
| CHRF_ENV_08 | *Klebsiella* | *K. quasipneumoniae* | *quasipneumoniae* | *K. quasipneumoniae* | *quasipneumoniae* |
| CHRF_ENV_09 | *Klebsiella* | *K. quasipneumoniae* | *quasipneumoniae* | *K. quasipneumoniae* | *quasipneumoniae* |
| CHRF_ENV_10 | *Klebsiella* | *K. quasipneumoniae* | *quasipneumoniae* | *K. quasipneumoniae* | *quasipneumoniae* |
| CHRF_ENV_11 | *Klebsiella* | *K. quasipneumoniae* | *similipneumoniae* | *K. quasipneumoniae* | *similipneumoniae* |
| CHRF_ENV_12 | *Klebsiella* | *K. quasipneumoniae* | *similipneumoniae* | *K. quasipneumoniae* | *similipneumoniae* |
| CHRF_ENV_13 | *Klebsiella* | *K. quasipneumoniae* | *similipneumoniae* | *K. quasipneumoniae* | *similipneumoniae* |
| CHRF_ENV_14 | *Klebsiella* | *K. variicola* |  | *K. variicola* |  |
| CHRF_ENV_15 | *Klebsiella* | *K. variicola* |  | *K. variicola* |  |
| CHRF_ENV_16 | *Klebsiella* | *K. variicola* |  | *K. variicola* |  |

### Analytical performance and cost of the assay

The final WGS-confirmed validation set of 61 KpSC and 4 non-KpSC isolates from clinical and environmental sources demonstrated concordance with MAMA PCR results with sensitivity of 100% (95% CI: 94 - 100) and specificity of 100% (95% CI: 94 - 100) (Figure 3C).

As a conventional agarose gel-based PCR assay, this approach requires only standard molecular laboratory infrastructure and is substantially more accessible than MALDI TOF mass spectrometry or whole genome sequencing. The estimated reagent cost per isolate was approximately USD$ 1 for the complete two reaction assay, supporting its potential use as a low-cost method for routine differentiation of KpSC members in resource limited settings.

## Discussion

We developed a conventional *rpoB*-based multiplex MAMA PCR for species- and subspecies-level differentiation of members of KpSC. The assay showed complete agreement with WGS for all experimentally tested targets and performed consistently across both clinical and environmental isolates. Using standard PCR and agarose gel electrophoresis, it offers a practical alternative for KpSC differentiation in laboratories without routine access to MALDI-TOF mass spectrometry or WGS. Given the importance of accurate KpSC identification for understanding disease burden and transmission dynamics (12), this assay provides a simple and accessible approach for routine laboratory use with a reagent cost of approximately $1 per sample.

A key advantage of the assay is its use of discriminatory SNPs in the conserved housekeeping gene *rpoB*. Analysis of 1,082 genomes showed that *rpoB* contains sufficient sequence variation to separate KpSC members at species and subspecies level, including discrimination of *K. quasipneumoniae* subsp. *quasipneumoniae* and subsp. *similipneumoniae*. Despite being a conserved housekeeping gene, *rpoB* showed clear phylogenetic clustering across the complex, enabling identification of SNPs that were consistently conserved within each taxon while remaining distinct across taxa. This provides an important practical advantage over existing low-complexity methods that either resolve fewer KpSC members, do not distinguish subspecies, or require real-time PCR platforms, which are more expensive to buy and maintain relative to conventional thermocyclers. In addition, this SNP-based strategy differs from earlier PCR-based approaches that rely on accessory or variably present targets, which may be more susceptible to horizontal gene transfer and therefore less stable for long-term differentiation (14, 15). Notably, the inclusion of primer sets for less commonly targeted members such as *K. africana* and *K. quasivariicola* broadens assay coverage beyond that of many currently available PCR-based methods (13, 17, 31, 32).

The validation results suggest that the selected *rpoB* SNPs are robust across diverse isolates. The assay was fully concordant with WGS for 49 clinical isolates representing 49 distinct STs across KpSC. This indicates that the selected *rpoB* SNPs are not restricted to specific lineages but remain robust across genetically diverse clinical backgrounds. In addition, 179 presumptive *Klebsiella* isolates recovered from wastewater samples were tested using the MAMA PCR, of which 174 were assigned to KpSC members, including *K. pneumoniae*, both subspecies of *K. quasipneumoniae*, and *K. variicola*. Representative environmental isolates were confirmed by WGS, showing complete agreement with PCR-based identification. Among the five isolates that produced inconclusive banding patterns, WGS of four isolates revealed that they were not KpSC members but *Enterobacter* spp., specifically *E. cloacae, E. roggenkampii*, and *E. hormaechei*. These findings are important because they show that the assay did not misclassify non-KpSC isolates as members of the complex. Instead, inconclusive amplification patterns correctly flagged them as outside the intended target group. Together, these results support the use of this assay as a confirmatory differentiation tool following routine microbiological identification of KpSc isolates and further highlights the limitations of phenotypic identification alone.

This study should be considered within the context of several limitations. First, the assay is intended for use on cultured isolates rather than directly on clinical specimens and still depends on prior conventional microbiological isolation and preliminary identification. Second, the workflow requires two PCR reactions to detect and differentiate all targeted KpSC members, which adds some operational complexity compared with single-reaction assays. Third, primers targeting *K. quasivariicola* and *K. africana* could not be experimentally validated on representative isolates because such strains were unavailable. In addition, *K. variicola* subsp. *tropica* was not included because its clinical relevance remains uncertain and genomic sequences from clinical sources are unavailable (33). Finally, while it costs $1 to conduct this assay in Bangladesh, associated costs might vary in different settings. Future work can include broader validation using larger and more geographically diverse isolate-collections, as well as direct comparison with existing molecular and non-molecular identification methods.

Overall, this *rpoB*-based multiplex MAMA PCR provides a simple, accurate, and low-cost method for differentiating common KpSC members in resource-limited settings. More broadly, the use of discriminatory SNPs within a conserved housekeeping gene may provide a useful framework for differentiation of other closely related bacterial complexes, such as *Burkholderia cepacia* complex (34) where accurate low-complexity identification remains challenging.

## Acknowledgments

We would like to thank Mr. Dipu Das, Ms. Sumona Akter, Ms. Deb Purna Keya, Ms. Esha Kazi, and Mr. Mohimenul Haque Rolin of the Child Health Research Foundation (CHRF), Bangladesh, for their help with microbiological and sequencing data generation, and bioinformatic analysis.

## Funding

The study was funded by Gates Foundation grant INV073135.

